# Power dynamics of theta oscillations during goal-directed navigation in freely moving humans: A mobile EEG-virtual reality T-maze study

**DOI:** 10.1101/2021.10.05.463245

**Authors:** Mei-Heng Lin, Omer Liran, Neeta Bauer, Travis E. Baker

## Abstract

Theta oscillations (∼4–12 Hz) are dynamically modulated by speed and direction in freely moving animals. However, due to the paucity of electrophysiological recordings of freely moving humans, this mechanism remains poorly understood. Here, we combined mobile-EEG with fully immersive virtual-reality to investigate theta dynamics in twenty-two healthy adults (aged 18–29 years old) freely navigating a T-maze to find rewards. Our results revealed three dynamic periods of theta modulation: 1) theta power increases coincided with the participants’ decision-making period; 2) theta power increased for fast and leftward trials as subjects approached the goal location; and 3) feedback onset evoked two phase-locked theta bursts over the right temporal and frontal-midline channels. These results suggest that recording scalp EEG in freely moving humans navigating a simple virtual T-maze can be utilized as a powerful translational model by which to map theta dynamics during “real-life” goal-directed behavior in both health and disease.

## Introduction

Decades of single-unit electrophysiological recordings of freely moving rodents navigating towards a selected goal (e.g. food, water, mates, shelter or avoiding danger) have produced a wealth of information about the neural mechanisms underlying goal-directed navigation (*1–4*). From this work, the consensus view is the precise firing rates of hippocampal place cells and parahippocampal grid cells with respect to the theta rhythm (4–12 Hz in rodents) constitute a temporal mechanism for encoding spatial position and information during navigation (*1*). In particular, theta oscillations have been shown to encode movement speed, direction, distance traveled, and proximity to spatial boundaries(*1, 5*). When salient events or cues such as rewards and navigationally-relevant landmarks are presented in the animal’s environment, the phase of the theta rhythm is reset, a process that appears to facilitate the encoding of salient information within the hippocampal-parahippocampal circuitry (*6*). Further, recent studies suggest that resetting the phase of the ongoing theta rhythm to endogenous or exogenous cues facilitates coordinated information transfer within hippocampal-parahippocampal circuits and between distributed brain areas involved in navigation (*7*). Computational work leverages such theta mechanisms to simulate the spatial distribution of firing fields of place and grid cells (*8, 9*). For example, computational models integrating spatial representations in the hippocampal-parahippocampal circuit explicitly require velocity-dependent modulation of theta oscillations (both frequency and power) in their contribution to path integration and navigation (*6, 10, 11*). Further, grid cell models require an input conveying the speed and direction of motion (i.e. velocity), information carried by theta rhythmicity, so that this spatial information can be integrated to estimate changes in location based on the distance and direction travelled(*8, 9*). Grid cells models also require phase-resetting of velocity dependent theta oscillations by location-specific input from place cells to prevent accumulation of error (*6, 8, 10*). Although theta dynamics during navigation have been well studied in non-human animal and computational work, whether theta oscillations are fundamental components of the brain’s navigation system in freely moving humans remains elusive.

This apparent lack of knowledge is likely due to the necessarily limited options for using invasive recording techniques in healthy humans subjects (*12*), and whilst animals can be examined during free movement, human studies employing virtual reality to simulate aspects of “real-world” navigation rarely achieve equivalent realism (*13*). Virtual reality can refer to one of three types of system: a virtual environment presented on a flat screen display (2D), a room-based system such as a CAVE, or a head-mounted VR display (3D). Traversing through any rendered environment via button presses or a joystick while physically immobile can result in motion sickness, sensory conflict, impair spatial navigation, and clearly influence the degree of immersion and presence in the virtual environment (*13, 14*). Notwithstanding, intracranial EEG recording in epilepsy patients have demonstrated the presence of movement-related theta oscillations in both the neocortex and hippocampus during immobile virtual navigation(*15, 16*). EEG and MEG studies have also identified functional parallels between theta oscillations (4-8 Hz in humans) recorded during immobile virtual navigation and those found in rodents during active navigation (e.g. self-initiated movement, processing of landmarks, path integration, orientation) (*17–23*). And two decades of fMRI studies have consistently demonstrated the involvement of several nodes of the navigation network (e.g. hippocampus, parahippocampal cortex, posterior parietal cortex, precuneus, the retrosplenial complex, and a region around the transverse occipital sulcus) during immobile virtual navigation tasks (*1, 2, 24–26*). Notably, Doeller et al. (2010) observed that the fMRI BOLD response in human right parahippocampal cortex exhibited a speed-modulated six-fold rotational symmetry in running direction as predicted by theoretical models of theta phase coding of grid cells(*27*).

While there is no doubt that the integration of neuroimaging and videogame design techniques have advanced our understanding of spatial navigation in humans, fMRI data lack the temporal and frequency information needed to study theta oscillations during navigation tasks(*28*), and immobile navigation lacks the self-motion information from visual, vestibular, proprioceptive and motor systems needed to generate the theta-dependent firing patterns of place and grid cells observed in rodent studies(*13, 29*). Thus, previous research has been unable to fully address whether freely moving humans also exhibit theta dynamics (e.g. phase-reset, movement speed and direction modulation) during mobile navigation. In recent years, several technological and methodological advances in electrophysiological research (mobile-EEG) and fully immersive virtual-reality (head mount display) have made mobile spatial navigation amenable for investigation in humans(*13, 14, 30*). Such investigation have already shown compelling results. For example, relative to standing still, delta-theta (2–7.21 Hz) power has been shown to increase during walking in an immersive virtual city (omnidirectional treadmill)(*30*) and in an virtual Y-maze housed in a large physical room(*14*), findings consistent with intra-hippocampus EEG recordings during real and virtual navigation(*31*).

Here, we leveraged this advancement to investigate theta dynamics in humans freely navigating a T-maze to find rewards. T-maze paradigms have been used extensively across several animal species (e.g. mice, rodents, ferrets, cats, squirrel monkeys, horses, cows, goats and sheep) to investigate “real-life” goal-directed navigation(*4, 32, 33*). The simplicity of the T-maze paradigm belies its utility and versatility for examining goal-directed navigation, and such investigations have produced a wealth of information about spatial learning and memory, reinforcement learning, and effort-based decision-making (*4, 32, 34–36*). Thus, the T-maze constitutes a natural application for mobile-EEG and immersive VR, providing a means for building a translational model of goal-directed navigation across species. Here, we recorded EEG from humans actively navigating a fully immersive virtual reality T-maze task to find rewards. Our purpose was 2-fold. First, given the novelty of the task, we wished to demonstrate that reward cues presented in the T-maze would evoke two well-established phase-locked theta responses, frontal-midline theta (FMT)(*37*) and right-posterior theta (RPT)(*17*). Second, in line with animal and computational work, we wish to demonstrate that the participant’s walking trajectory (leftward vs rightward trials) and speed (fast vs slow trials) towards the feedback location would differently modulate theta activity. Taken together, these results provide converging evidence for the proposal that task and behavioral variables (reward, direction, and speed) are responsible for modulating theta activity during active navigation, and hold out promise for integrating experimental, computational, and theoretical analyses of goal-directed navigation in animals within the field of human EEG research.

## Results

### Behavior

In this study, twenty-two young adults (20 right-handed [laterality index = 68], 9 male and 13 female, aged 18–29 years old [M = 21, SE =.61]) freely navigated a T-maze to find rewards (Fig. 1A). On average, participants completed 148 trials (SE = 7.03, range = 100 – 238), and took 4.2 seconds (SE = .14, range = 2.97 – 5.63) to reach the feedback location (1.83 m). Overall, no differences were observed between the percentage of leftward (M = 48%, SE = 4.3) and rightward (M = 52%, SE = 3.6) trajectories, t(21) = −1.6, p = .123, nor their velocity (leftward: M = .449 m/s, SE = .015 | rightward: M = .448 m/s, SE = .016) towards the feedback location, t(21) = .364, p = .719. It is worth noting that participant first 15 trials were biased towards rightward turns, t(21) = 2.6, p < .01. In regards to post-feedback behavior, participants adopted a Lose-shift strategy (M = 71%, SE = 3.28), t(21) = −6.05, p < .001, and were faster for Win-stay trials (M = .46 m/s, SE = .017) relative to Win-shift trials (M = .44 m/s, SE = .016), t(21) = 3.7, p < .001.

**Figure 1.**
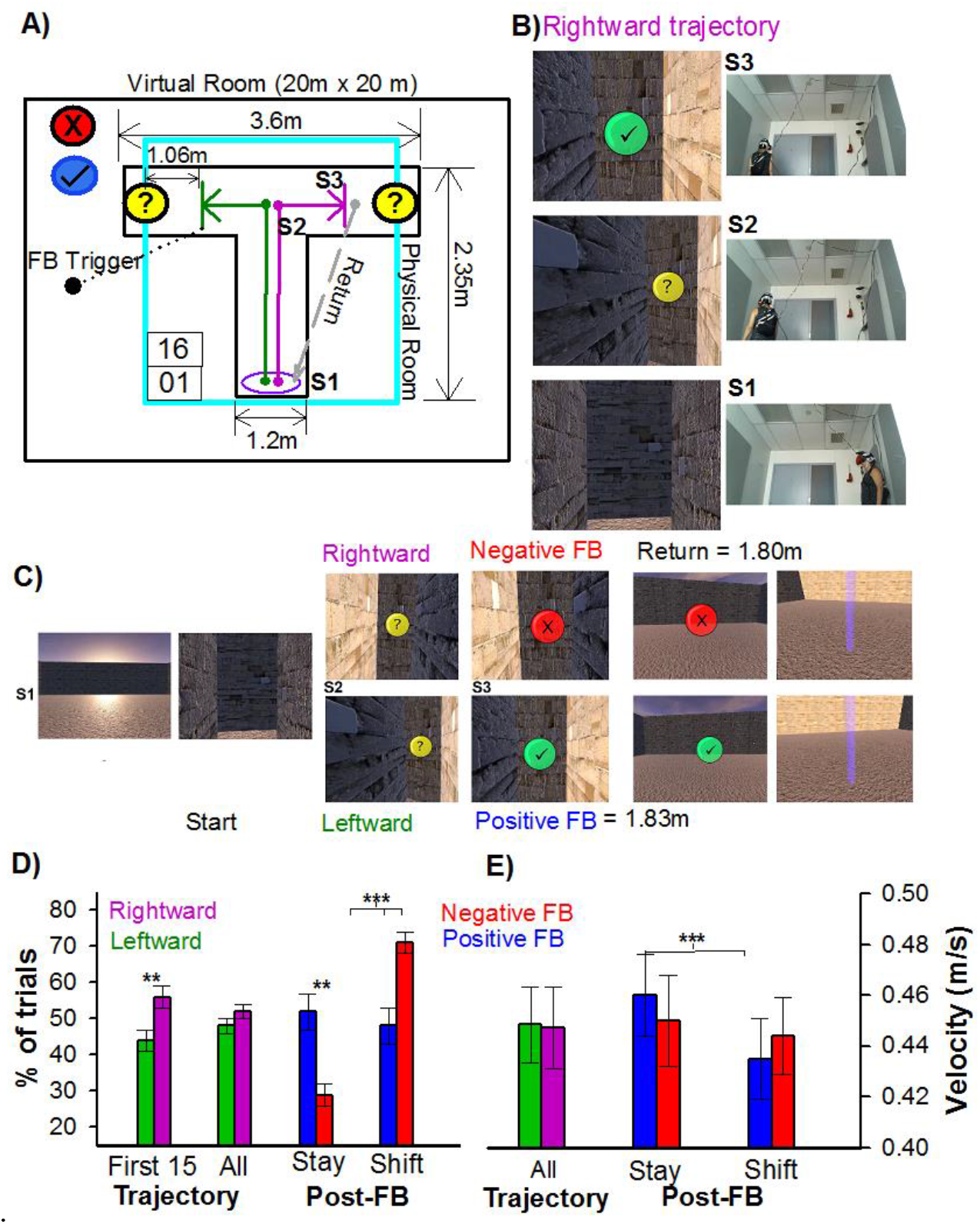
Mobile virtual reality T-maze paradigm and associated behavior. **A)** Dimensions of the virtual (black border) and physical (cyan border) room and T-maze (S1: start location, S2: junction point, S3: feedback location). Purple and green lines denotes rightward and leftward trajectories, respectively. **B)** An example of a rightward trajectory in the T-maze, **(C)** and trial-to-trial sequence of events. Behavioral analysis for choice **(D)** and velocity **(E).** Green and purple bars denote leftward and rightward trajectories, and Blue (positive) and Red (negative) bars denote post-feedback behavior.

### Feedback-related Theta Responses

Given the novelty of the mobile-EEG T-maze paradigm, sa necessary precursor would be to replicate two well-studied feedback-related EEG responses observed using conventional computer-based 2D tasks, frontal-midline theta (FMT)(*37*) and right-posterior theta (RPT)(*17, 26*). FMT describes an obligatory pattern of phase reset and power enhancement in frontal-midline electrodes (4–8 Hz: 220–300 msec) found to be sensitive to the valence of the feedback (e.g. increase in power and phase consistency following negative feedback), and has been associated with midcingulate cortex processes related to cognitive control and reinforcement learning (*37, 38*). This phenomenon is also observed in the time domain as a component of the event-related brain potential (ERP), called the feedback-related negativity or N200. RPT describes a pattern of phase reset and power enhancement in right-posterior electrodes (4–8 Hz: 160-220 msec) found to be sensitive to the spatial position of the feedback (e.g. greater power and phase consistency for feedback found following rightward turns relative to leftward turns), and associated with parahippocampal processes related to spatial navigation (*17, 26, 39, 40*). This phenomenon is also observed in the time domain as an ERP component called the topographical N170. To examine these two oscillatory components, we computed a standard single trial wavelet-based time-frequency analysis to the EEG signal time-locked to the onset of positive and negative feedback (FMT) following leftward and rightward turns (RPT).

Visual inspection of Fig. 2 reveals a clear enhancement of FMT power between 220 and 260 ms (peak power: M = 250 msec, SE = ±.14) and RPT power between 180 and 220 ms (peak power: M = 211 msec, SE = ±.14) following the onset of feedback stimulus. In regards to FMT, a repeated measures ANOVA on mean band power measured at Fz as function of Frequency (delta, theta, alpha, beta) and Valence (positive vs negative feedback) revealed a main effect of Frequency (F_(3, 63)_ = 19.67, p < .001, η_p_^2^= .48), and Valence (F_(1, 21)_ = 5.13, p < .05, η_p_^2^= .20), and an interaction between Frequency and Valence, F_(3, 63)_ = 3.31, p < .05, η_p_^2^= .144. Post-hoc analysis indicated that the EEG was characterized by greater power in the theta band (FMT, M = .30 dB, SE = ±.05) than at each of the other frequency bands (p < .01), and FMT power was greater for negative feedback (M = .36 dB, SE = ±.06) relative to positive feedback (M = .24 dB, SE = ±.04), t(21) = −2.3, p < .05, Cohen’s d = .52 (See Figure 2A). No other frequency bands displayed power differences between positive and negative feedback (p > .05). In regards to RPT, a repeated measures ANOVA on mean band power measured at P8 as function of Frequency (delta, theta, alpha, beta) and Trajectory (leftward vs rightward) revealed a main effect of Frequency, F_(3, 63)_ = 22.82, p < .001, η_p_^2^= .50, indicating that the EEG was characterized by greater power in the theta (M = .57 dB, SE = ±.09) and alpha (M = .44 dB, SE = ±.09) band than at each of the other frequency bands (p<.001). However, no other main effects nor an interaction were detected (p>.05). Together, these results are characteristic of FMT and RPT, and indicate that the feedback processing in the virtual reality T-maze task is capable of eliciting these phased-locked theta responses during active navigation.

**Figure 2.**
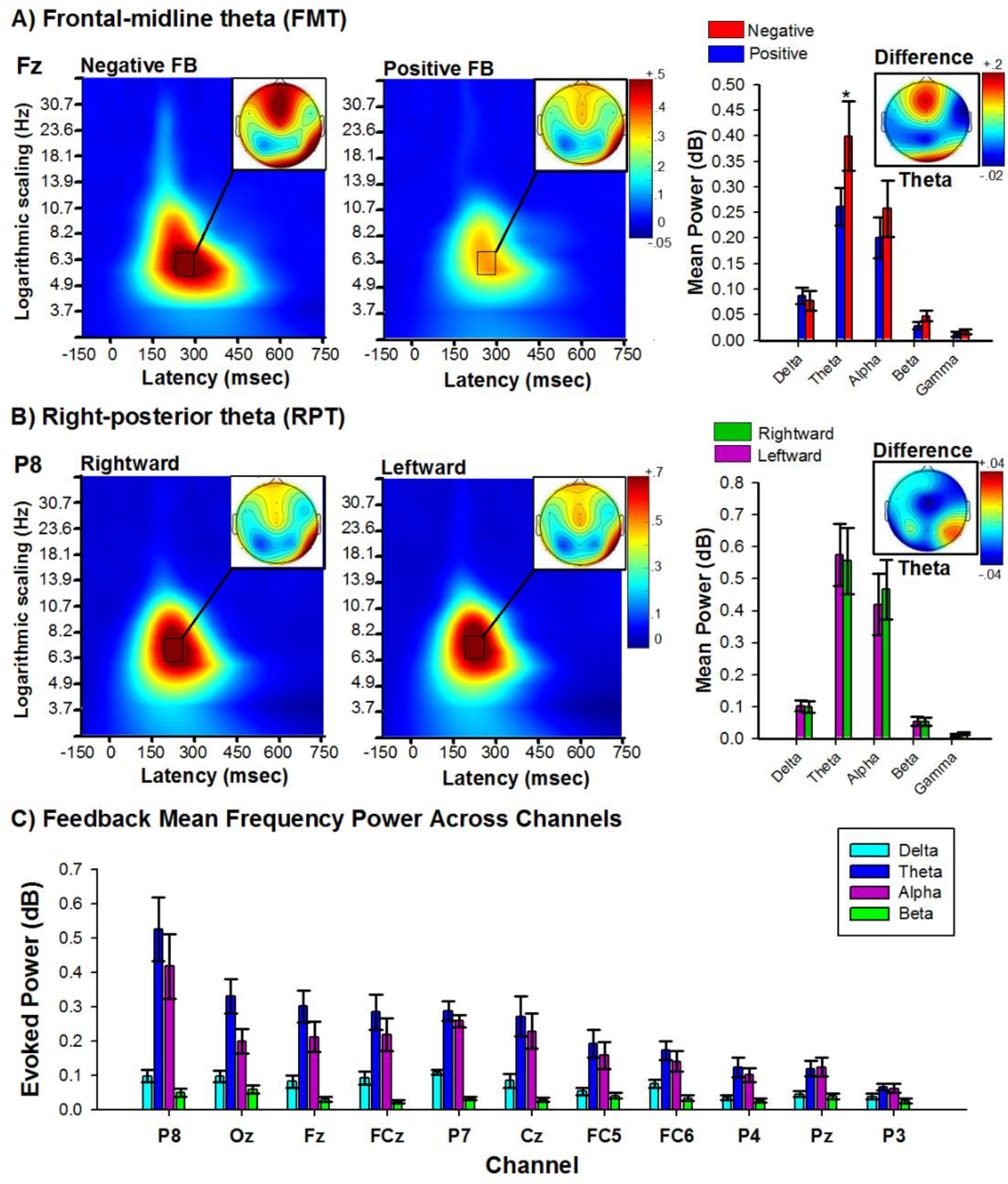
Feedback processing during active navigation. **A)** Panels indicate changes in power for each frequency band with respect to baseline (−300 to −100 ms period prior to feedback stimulus) elicited by negative (left) and positive (right) feedback stimuli. Right bar graph depicts peak power across frequency bands delta [1–3 Hz], theta [4–8 Hz], alpha [8–13 Hz], [13–20 Hz], and gamma [20–40 Hz] associated with the response to negative (red bars) and positive (Blue bars) feedback. Note highest power in the theta band, and stronger for negative feedback. Data recorded at channel Fz. **(B)** Panels indicate changes in power for each frequency band with respect to baseline (−300 to −100 ms period prior to feedback stimulus) elicited by feedback stimuli presented in the right alley (left) and in the left alley (right). Right bar graph. Peak power across frequency bands associated with the response to feedback in left (green bars) and right (purple bars) alley. Note highest power in the theta band, for both left and right alleys. Data recorded at channel P8. C). Bar graph illustrates the mean feedback power (150 – 300 ms) across frequency bands delta [1–3 Hz], theta [4–8 Hz], alpha [9–12 Hz], and beta [13–20 Hz] evaluated at all electrode channels, ordered by size. Bars indicate the standard error of the mean. Note highest power was in the theta band, and this increase in power exhibited a maximal at right posterior (channel P8). Error bars indicate the standard error of the mean.

### Movement-related Theta Responses

In line with animal and computational work, which have demonstrated that theta oscillations encode movement speed and direction during navigation, we examined whether the participant’s trajectory (leftward vs rightward trials) and walking speed (fast vs slow trials) towards the feedback location would differently modulate theta activity. We used the subjects median RT from start to feedback onset (distance travelled: 1.83 meters) to create two speed-dependent conditions (e.g., fast [median = .385 m/s, SE = .019] vs slow [median = .520 m/s, SE = .011]) and two direction-dependent conditions (e.g., leftward vs rightward trajectories). Fig. 3 illustrates the topography results of the time frequency analysis from the start location to the feedback location averaged across all conditions. Visual inspection of Fig. 3 reveals notable enhancements of theta power (as well as delta power) over frontal-midline (FCz and Cz) and posterior (P3, Pz) channels while traversing the stem (S1a and S1b) and turn (S2a and S2b) sections of the T-maze. We confined the statistical comparisons of the time-frequency space to these frontal and posterior electrodes (see Fig. 4). We also included an analysis of P8 because of its robust theta responses during feedback processing in the maze (Fig. 2B and 2C). Statistical comparisons of data for each grand averaged time-frequency plot were calculated using paired-samples t-tests (left vs right; fast vs slow). Given the large search space and novelty of this experiment, the alpha value was set at p < .05 (uncorrected) for each t-test conducted. To provide partial control for Type I error inflation, at least two consecutive significant comparisons (2 Bins of time data [approx. 50-100 ms] across two frequency steps [2 hz]) were required before a specific value was portrayed on the graph (*41*). This value was chosen as it provided the best visual representation of the differences between the conditions of interest, and a necessary precursor if we are to begin developing empirically driven and realistic representations of the oscillatory dynamics used to encode, represent, and process information during active navigation.

**Figure 3.**
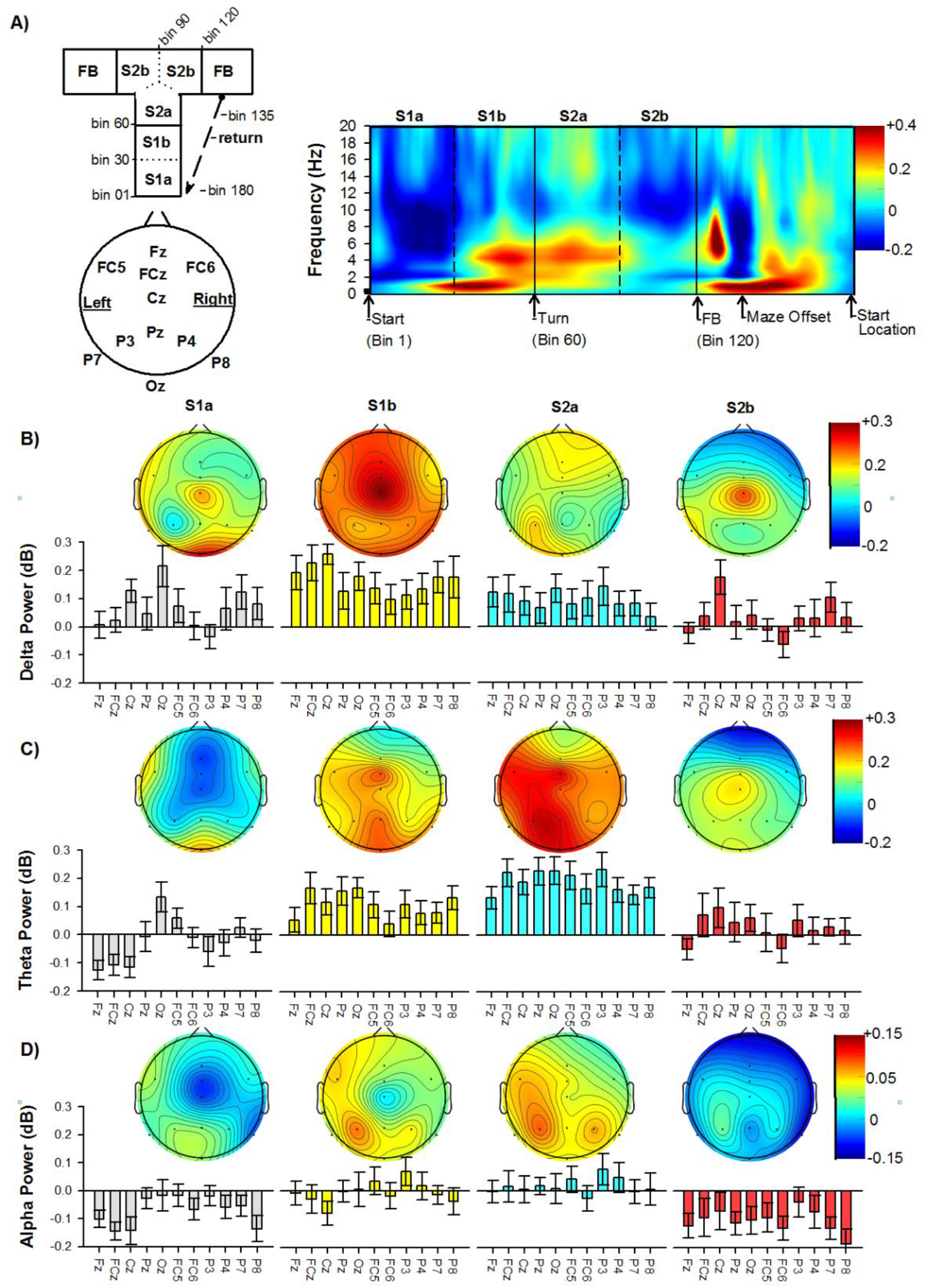
Frequency power and topography across the T-maze traversal. **A)** Top-left panel. A diagram illustrating the maze subsections and their associated Bin range. Bottom-left panel depicts the channel locations. Right panel indicate changes in power for each frequency band (with respect to baseline) averaged across all conditions and subjects at FCz. Topographical maps representing the mean frequency power at each channel for **B)** delta [1–3 Hz], **C)** theta [4–8 Hz], and **D)** alpha [9–12 Hz] for each subsection (S1a, S1b, S2a, S2b) of the path from the start to feedback location. Bars indicate the standard error of the mean.

**Figure 4.**
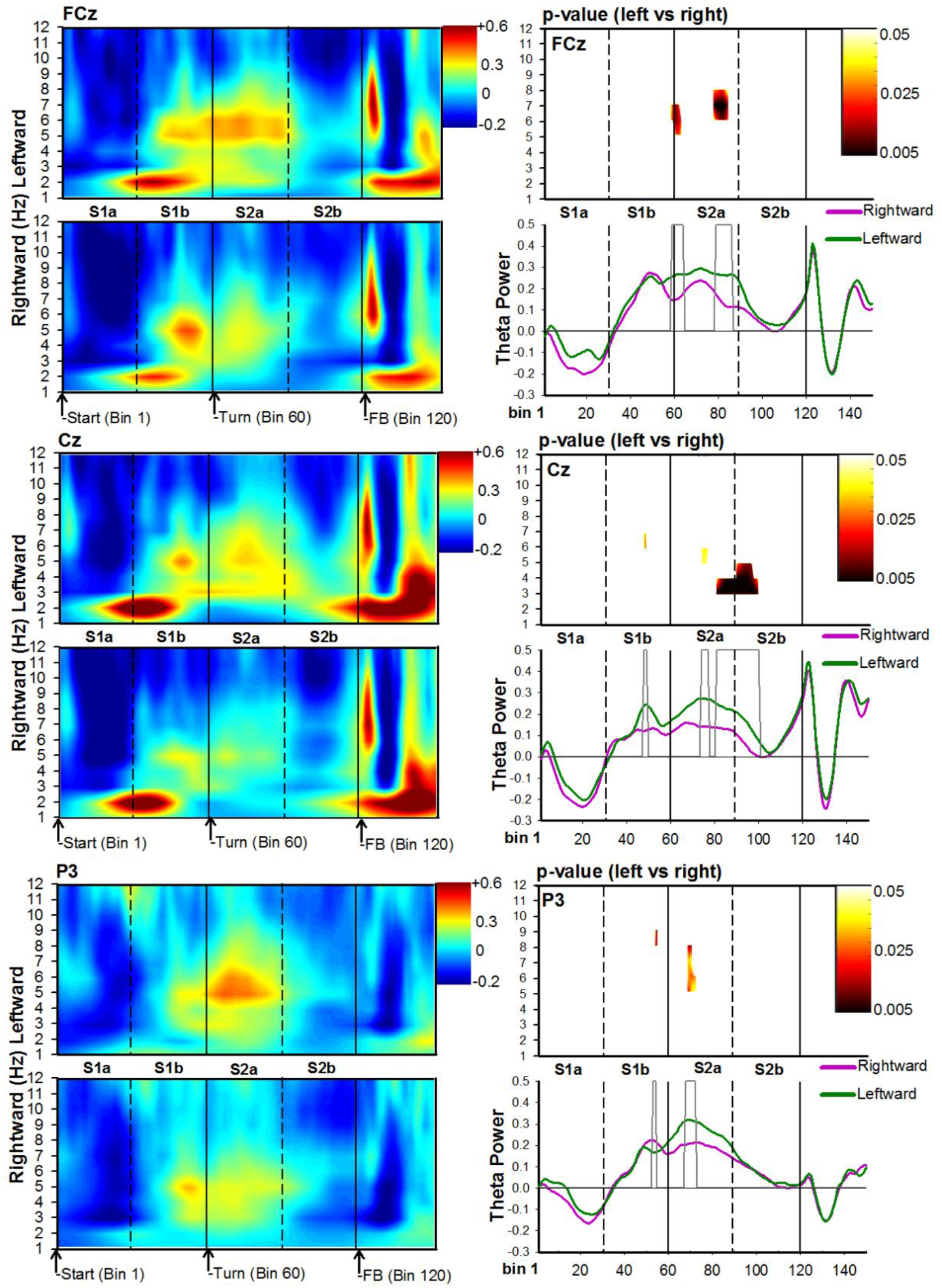
Time-frequency analysis associated with maze trajectories. For each channel location, FCz (top), Cz (middle), and P3 (bottom), panels depict time-frequency power maps (left panels), p-value maps (right-top panel), and theta time-course (right-bottom panel) for the leftward (green solid lines) and rightward (purple solid lines) conditions. The X-axis represents Bin location and maze subregion. The Y-axis for power and p-value maps represents frequency ranges from 0 to 12 Hz, and the Y-axis for the theta time-course represents a change in power. For all conditions, Bin 0 represents the start of the trial. The color bar for time-frequency plots represents the power of the oscillations depicting greater activity in warm colors. The heat-maps (left-top panel) represents the p-values (range .05 to .005) comparing leftward vs rightward trajectories. In particular, paired comparisons of data used to generate each grand averaged heat-map were calculated using paired-samples t-tests. The alpha value was set at .05 for each t-test conducted. However, to provide partial control for Type I error inflation, at least two consecutive significant comparisons around the target value were required before a specific value was portrayed on the graph(*41*). The grey bars depicted in the theta-time course maps represent significant Bins identified in the heat-maps.

To help visualize the subject’s location during their trajectory, we segmented the stem (S1a [Bin 1-30]; S1b [Bin 31-60]) and turn (S2a [Bin 61-90]; S2b [91-120]) sections of the T-maze (see Figure 3A). In regards to direction travelled, the first theta burst (6-7 Hz; channel Cz) occurred as participant approached the junction region of the T-maze (S1b Bin 48-49; duration = 84 msec), and displayed a sensitivity to leftward relative to rightward trajectories (range: t(21) = 2.2 – 2.7, p = .04 – .02). Channel FCz also displayed a similar pattern of results in the stem, but closer to the junction point of the maze (6-8Hz, Bin 59-64; duration = 192 ms; range: t(21) = 2.1 – 2.7, p = .05 – .01). As subjects arrived at the junction point (S2a), a second burst of theta could be seen across several channels, all of which maintaining a leftward sensitivity: Channel P3 (5-8Hz, Bin 68-72; duration = 135 ms; range: t(21) = 2.1 – 2.8, p = .05 – .01); Channel Cz (5-6Hz, Bin 74-77; duration = 108 ms; range: t(21) = 2.1 – 2.2, p = .05 – .03); and, Channel FCz (6-8Hz, Bin 79-86; duration = 210 ms; range: t(21) = 2.1 – 3.4, p = .05 – .002). As the subjects began their approach towards the goal location (S2a-S2b), there was a strong increase in delta-theta power at channel Cz for the left alley relative to the right alley: (3-5Hz, Bin 81-100; duration = 567 ms; range: t(21) = 2.2 – 4.8, p = .03 to < .00001). In addition, theta-alpha activity (8-9 hz) at channel P3 displayed a sensitivity to rightward trajectories (8-9Hz, Bin 53-54; duration = 84 ms; range: t(21) = 2.2 – 4.8, p = .02 – .009). It is also worth noting that the effects observed over channel P3 where not observed over channel P4 (channel P4 did not display any significant Bins for any frequency). Together, these results indicate that theta power was sensitive to the participant’s trajectory from the start location to the goal location in the T-maze.

In regards to speed (Figure 5 and 6), there was an initial increase in delta-theta power at the beginning of the stem, which was stronger for slow trials relative to fast trials at channel P8: (3-4Hz, Bin 39-40; duration = 84 ms; range: t(21) = 2.2 – 3.0, p = .04 – .007). Shortly after this response (approx. 800 ms), a theta burst emerged as the participant approached the junction region, and was stronger for slow trials: Channel FCz (5-6 Hz, Bin 51-54; duration = 168 msec; t(21) = 2.1 – 2.2, p = .04 – .02), and Channel P3 (6-7 Hz, Bin 56-59; duration = 168 ms; range: t(21) = 2.1 – 3.1, p = .04 – .006). By contrast, as subjects approached the goal location after the turn, a second burst of theta could be seen across several channels and all displayed an increase in power for fast trials: Channel FCz (7-8Hz, Bin 79-81; duration = 81 ms; range: t(21) = −2.1 – −3.5, p = .03 – .001); Channel Cz (4-7Hz, Bin 86-93; duration = 216 ms; range: t(21) = −2.1 – −3.2, p = .04 – .004); and, Channel P3 (4-6Hz, Bin 83-95; duration = 315 ms; range: t(21) = −2.2 – −3.1, p = .04 – .004). Together, these results indicate that theta power was also sensitive to the participant’s speed in the T-maze, but was stronger for slow trials as participants approached the junction point, and stronger for fast trials as participants approached the goal location.

**Figure 5.**
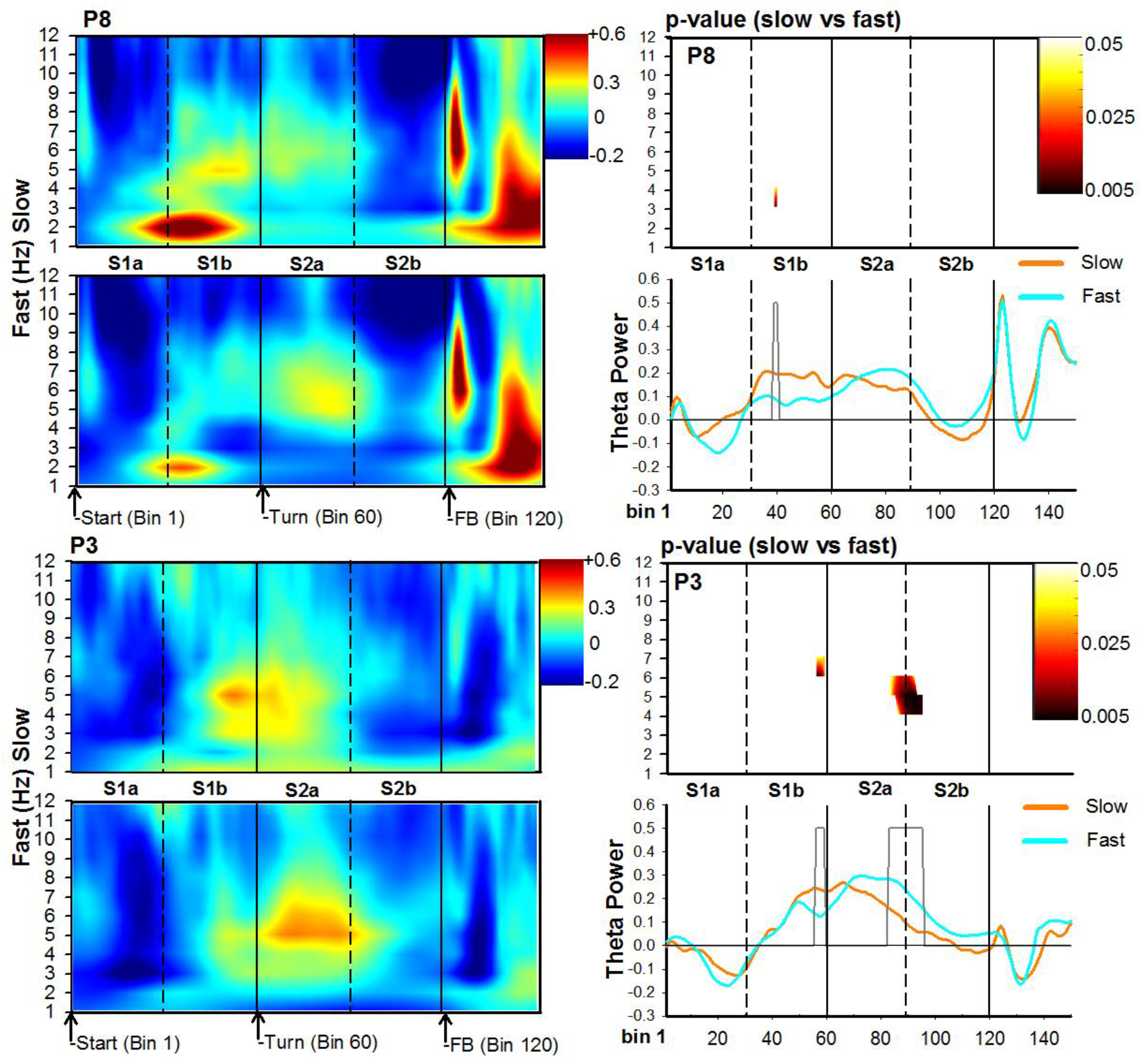
Time-frequency analysis associated with trial walking speed for posterior channels P8 (top) and P3 (bottom). For each channel location, panels depict time-frequency power maps (left panels), p-value maps (right-top panel), and theta time-course (right-bottom panel) for the slow (orange solid lines) and fast (cyan solid lines) conditions. The X-axis represents Bin location and maze subregion. The Y-axis for power and p-value maps represents frequency ranges from 0 to 12 Hz, and the Y-axis for the theta time-course represents a change in power. For all conditions, Bin 0 represents the start of the trial. The color bar for time-frequency plots represents the power of the oscillations depicting greater activity in warm colors. The heat-maps (left-top panel) represents the p-values (range .05 to .005) comparing slow vs fast trials. The grey bars depicted in the theta-time course maps represent significant Bins identified in the heat-maps.

**Figure 6.**
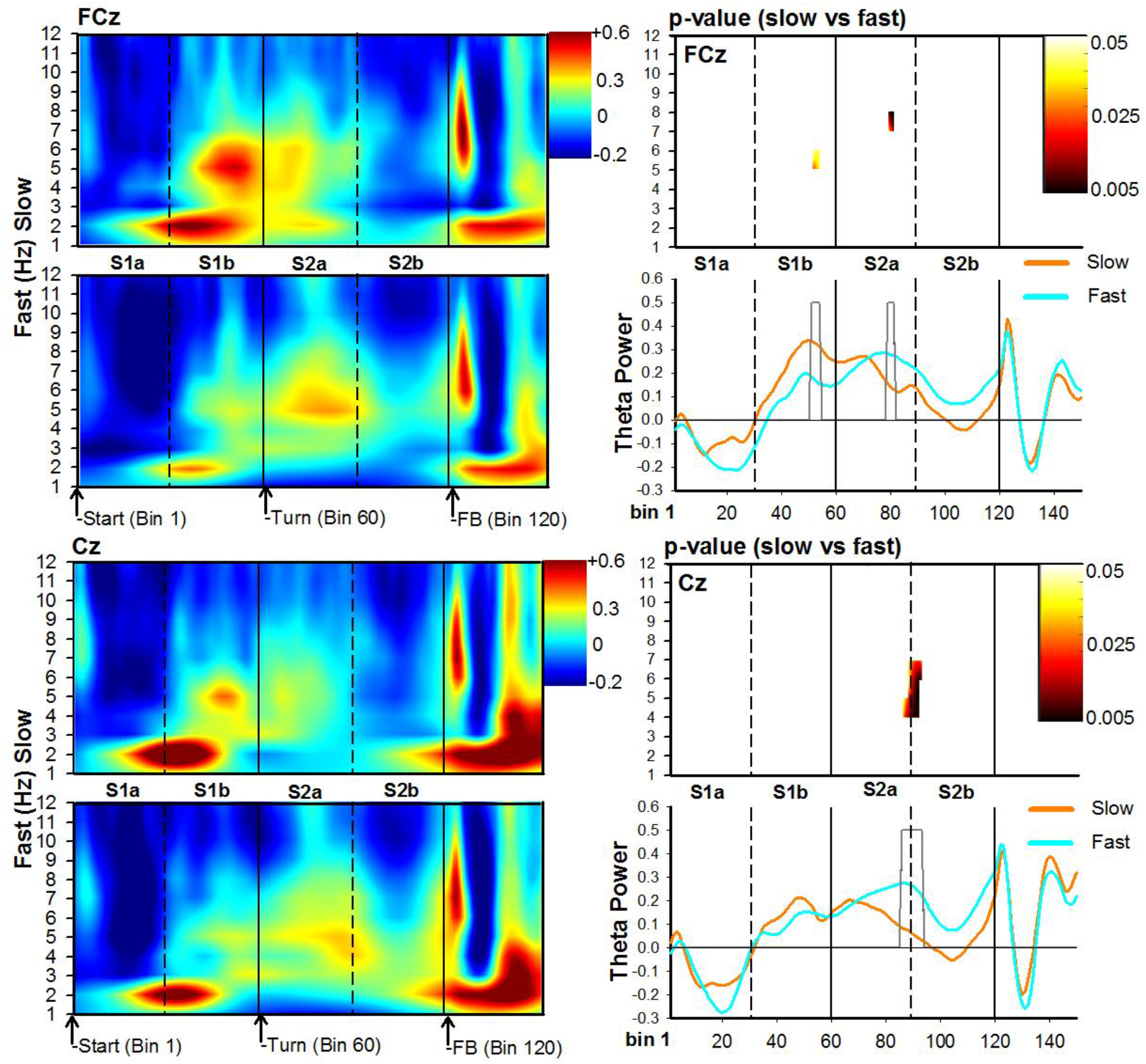
Time-frequency analysis of the EEG associated with trial walking speed for frontal-midline channels FCz (top) and Cz (bottom). For each channel location, panels depict time-frequency power maps (left panels), p-value maps (right-top panel), and theta time-course (right-bottom panel) for the slow (orange solid lines) and fast (cyan solid lines) conditions. The X-axis represents Bin location and maze subregion. The Y-axis for power and p-value maps represents frequency ranges from 0 to 12 Hz, and the Y-axis for the theta time-course represents a change in power. For all conditions, Bin 0 represents the start of the trial. The color bar for time-frequency plots represents the power of the oscillations depicting greater activity in warm colors. The heat-maps (left-top panel) represents the p-values (range .05 to .005) comparing slow vs fast trials. The grey bars depicted in the theta-time course maps represent significant Bins identified in the heat-maps.

## Discussion

In the present study, we combined mobile-EEG and head-mounted VR technology to investigate whether behavior (direction and speed) and task (rewards) variables modulate scalp-recorded theta activity in humans freely navigating a T-maze task. In line with animal and computational work, our results provide compelling evidence that theta power was dynamically modulated as participants traversed the T-maze towards the goal location and received reward feedback. Previous research in rodents, non-human primates, and humans suggests that at least three types of theta oscillations exist during navigation: one elicited during movement in space(*1*), another in response to planning and decision-making(*42*), and a third in response to reward processing(*37*). Our findings suggest that such theta-related responses were expressed across time and topography during the traversal of the T-maze.

### The Stem

Shortly after participants began their movement down the stem of the T-maze, a large increase in delta power was observed over the right medial temporal (P8) and frontal-midline (Cz) electrodes. Prior rodent and human studies have also revealed similar patterns of movement-related increases in delta activity (*15, 30, 43, 44*). For example, EEG studies using joystick-based movements through 2D rendered virtual environments suggest that movement-related oscillations based on optic flow tend to manifest specifically within the 1–8 Hz frequency range (*31, 43*). More recently, Liang and colleagues (2018) demonstrated that frontal-midline delta-theta oscillations (2–7.21 Hz) exhibit higher power and are more sustained during physical movement than when standing still on an omnidirectional treadmill coupled with 3D immersive virtual reality. Delaux et al. (2021) also observed greater delta power as participants began walking down the starting arm of a fully immersive 3D Y-maze. Together, these data suggest that delta-theta oscillations can be induced by movement via a combination of visual, vestibular, and proprioceptive information. Further, while this emerging pattern of delta activity advocates for a mere signature of locomotion, its worth noting that delta-theta (3-4 Hz) activity recorded over right medial temporal cortex (electrode P8) proved to be condition sensitive, i.e., higher power for slow walking trajectories relative to fast walking trajectories. Consistent with this finding, Delaux et al. reported stronger delta response during learning phases of their Y-maze task, and intra-hippocampus EEG recordings found a delta-theta sensitivity to different types of real-world movements (e.g. during searching, recall and walking) during real and virtual navigation (*31*). Further, several human studies suggest that virtual navigation tends to result in low-frequency hippocampal oscillations peaking around 3.3 Hz, whereas freely ambulating humans show increased hippocampal oscillations ranging from 1–12 Hz compared with a standing position (*15, 31, 43, 45*). Although parallels exist between scalp recorded EEG and intracranial EEG recordings, the hippocampus is located too deep in the brain to be detected with electrodes placed at the scalp and because of its spiral organization, would likely produce a closed electromagnetic field (*17, 40*). This concern notwithstanding, movement-related signals conveyed by the hippocampus project to and regulate navigation regions in temporal, parietal, and prefrontal cortex (*15, 23, 46*), and these regions are amenable to investigate with scalp EEG(*28, 47*). Thus, the movement-related delta-theta activity observed here, and in other mobile EEG-VR studies, may be a cortical reflection of the movement-specific firing patterns of the hippocampal circuitry observed in intracranial EEG studies, and highlight the importance of ambulation to the induction of low-frequency oscillations and to spatial processing(*13, 29*).

### The Junction

As participants approached the junction section of the T-maze, a burst of frontal-midline theta power emerged and exhibited an increase in power for slow and leftward trajectories. Although this theta response deviates from previous observations of proportional increases in delta/theta activity with increases in velocity, it’s worth noting that this increase in theta power coincided with the participants’ decision-making period, and before the turning motion itself. For these reasons, we propose this increase in frontal-midline theta activity may be more in line with route planning and decision-making. In particular, when animals come to a decision point in a T-maze, they sometimes pause or slow down as if deliberating over the choice (i.e. mentally searching future trajectories) (*42*). Neurophysiological data in rodents suggest that increases in hippocampal place cell activity during this period represent the process in which the animal is serially exploring the paths towards future outcomes (*42, 48*). Several researchers have further suggested that coherent oscillations between prefrontal cortex and hippocampus create such imagined episodic futures for this purpose (*42, 49, 50*). Further, hippocampal theta-entrainment of the rodent medial prefrontal cortex is strongest near the decision-making period of spatial memory tasks, which serves to focus attention on the prefrontal representations that are relevant for task performance (*51–54*). For example, a previous study revealed increased theta-entrainment between medial prefrontal and hippocampal neurons at the choice point of a working memory T-maze task (*55*). In humans, deliberative decision-making is also hypothesized to involve the prefrontal cortex and medial temporal lobe structures, suggesting that there are direct parallels between animal and human findings(*42*). For instance, neuroimaging evidence revealed that the hippocampus is both necessary for and active during episodic future thinking(*56*), and several EEG studies have also shown that when subjects engage in control processes characterized by goal-directed influence, there is an increase in frontal theta activity (*7, 37, 57–59*). Together, these studies highlight the role of hippocampal-prefrontal theta interactions across different cognitive domains, such as goal-directed behavior(*7*), episodic memory (*23*), decision-making (*42*) and spatial learning (*52*). By extension, we propose that the observed increase in right posterior delta-theta power and frontal-midline theta power during slow trials may dovetail the neural processes and theoretical assumptions of deliberative decision-making observed across species. These findings imply that when reward-delivery contingencies are variable, humans at decision points in a T-maze, like rodents, are actually searching through possibilities, evaluating those possibilities, and making decisions that are based on those evaluations, a process reflected by an increase in both response time (i.e. slowing or pausing) and the presence of temporal-frontal theta oscillations near decision points(*42*), as we observed here.

Moreover, we propose that the observed increase in frontal-midline theta power for leftward trajectories may reflect additional control processes by frontal cortex during the decision-making period. Studies in rodents, non-human primates, and humans have uncovered signals in the anterior midcingulate cortex that reflect the pressure to switch away from an ongoing behavioral strategy or default action (*60*). Frontal-midline theta activities, which are proposed to be generated in anterior midcingulate cortex(*37*), have also been shown to predict behavioral switching in simple reinforcement learning tasks(*38*), and are enhanced during more cognitively demanding navigation periods in spatial tasks (*18, 19, 57*). In parallel, since the 1920s preferences in turning direction have been reported in several animal species, including humans(*61, 62*). For instance, a rightward turning bias in humans can be observed when walking around obstacles or making turns in a T-maze(*62*). Consistent with this turning bias, 65% of participants in the present study displayed a rightward turning bias at the beginning stages of the task, possibly reflecting the default action in the T-maze. In consideration of these observations, we propose that the increase in frontal-midline theta power prior to the junction point of the T-maze may reflect anterior midcingulate cortex control response to switch from the default action of turning right, to the non-preferred action of turning left. In other words, the observed increase in frontal-midline theta activity reflects the increased switch demand by anterior midcingulate cortex that would be required to implement top-down control across disparate brain regions to override the tendency to turn right. Although admittedly speculative, we hope these findings will motivate future experimental and theoretical analysis of the neural determinants of human behavior at a choice-point in a T-maze.

### The turn and goal approach

From the junction point throughout the traversal of the turning section of the maze, the increase in frontal-midline theta power for leftward trials was sustained, possibly reflecting the maintenance period of the selected leftward action. Consistent with this observation, a previous mobile virtual reality study demonstrated a sustained theta response from the center zone of a Y-maze to the finish arm (*14*). We propose that this sustained frontal-midline theta response is likely generated by prefrontal cortex (e.g. anterior midcingulate cortex). According to an influential learning theory of anterior midcingulate cortex function, this region not only selects sequences of actions during the decision making process, but also determines the level of effort to be applied toward executing the action and maintaining this level of activity until the organism reaches its goal(*63*). Consistent with this view, a multitude of studies have indicated that frontal-midline theta power correlates positively with levels of cognitive effort, working memory load and attention, especially for tasks that demand sustained effort and control(*37, 64*). Based on this theoretical and empirical work, we propose that the frontal-midline theta activity observed following the junction point represents the continued engagement of the anterior midcingulate cortex and it’s role in maintaining vigilance and control of the leftward trajectory towards the goal location.

Following the junction point, leftward trajectories towards the goal location produced a strong theta burst over the left posterior channel P3. To note, this pattern of theta activity (or the inverse of) was not observed over the right posterior channel P4, ruling out the possibility that this enhancement of power was related to head-direction, motion artifacts, or stemmed from a hemispheric bias associated with the retinotopic position of the goal target stimuli (floating orb) during the turn. While the topography of this theta response was not anticipated, the robustness of its effects warrants a closer look. Based on the literature and topography of this theta response, one possible generator is the posterior parietal cortex(*65*). A large number of studies across species have related posterior parietal cortex activity to the control of body movements (e.g. eyes, head, limbs, and body), decision-making, and spatial navigation (*66–72*). In particular, posterior parietal cortex firing patterns in rodents are often determined by conjunctions of body position or orientation, positions in a path, and concurrent movement type (i.e., turns or forward locomotion)(*68, 73, 74*). For example, Krumin and colleagues (2018) trained mice to use vision to make decisions while navigating a virtual reality task, and found that posterior parietal cortex activity can be accurately predicted based on the position of the animal along the corridor and heading angle. These data, along with others, have led to the idea that posterior parietal cortex activity form an integration of spatial representations of objects and scenes with motor representations to support accurate eye, head, and whole body movements towards selected goal or target (*69, 75*). Relevant to motor coordination during the pursuit of goals, posterior parietal cortex activity also exhibits a sensitivity to self-motion (e.g. linear and angular speed), visual target position, and movement direction in egocentric coordinates. These findings help support the idea that posterior parietal cortex may subserve online sensorimotor coordination necessary for goal pursuit behavior or target chasing in egocentric space(*76*). By extension, we propose the theta activity recorded over the left parietal cortex during the turn may reflect the sensorimotor coordination process of pursuit navigation, (i.e., the continuous adjustment of movement plans relative to the position of the floating goal orb in the left or right alley of the T-maze). Further, the heightened activity for leftward trajectories likely represents the allocation of top-down control by anterior midcingulate cortex over posterior parietal cortex activity during the active pursuit of the leftward goal. We hope these findings will warrant future investigations.

Lastly, an increase in theta power over frontal-midline (FCz and Cz) and left posterior (P3) electrodes was observed during fast walking trajectories towards the goal target, findings consistent with previous observations of proportional increases in theta activity with increases in speed. In particular, animal and computational work indicate that theta oscillations coordinate the firing patterns of hippocampal place cells and parahippocampal grid cells during navigation, providing the rodents spatial position in the environment(*1, 6, 11*). Central to this idea is the observation that the power (and frequency) of hippocampal and parahippocampal theta activity increases linearly with movement speed, and such speed-related changes in theta oscillations is essential to calculate the distance travelled through the place field(*20, 44, 77*). Speed-related changes in theta power have also been linked to changes in sensorimotor integration, the flow of sensory input, as well as cognitive/memory functions(*44*). For instance, the sensorimotor integration hypothesis posits that rodent hippocampal theta oscillations incorporate incoming sensory information with existing motor plans to guide movement, and more rapid traversals require faster sensorimotor transformations, resulting in higher theta activity(*20, 57*). Regardless of the theoretical interpretation of speed-related changes in theta power during navigation, the observed speed- and direction-related increase in theta power during the approach to the goal location draw strong parallels with animal and computational studies. Further, although these specialized neural representations have been identified in humans during virtual movement at various levels of analysis - i.e., ranging from intracranial EEG recordings of local field potentials to the fMRI blood oxygen level-dependent (BOLD) signal - virtual movement and real movement are fundamentally different(*13*). Virtual movement requires subjects to press buttons or move a joystick to process optic flow in order to compute their speed, direction, and location in space, and to initiate and maintain virtual movement toward the target location, all while physically immobile(*13, 14, 30*). By contrast, self-motion information from visual, vestibular, proprioceptive and motor systems are needed to generate the theta-dependent firing patterns of hippocampal-parahippocampal system. Thus, our findings here confirms that spatial navigation and free ambulation are potential drivers of multiple theta generators in healthy human participants, and likely reflects the common theta state the navigation system is synchronized to(*15*). More specifically, given the role of hippocampal theta in synchronizing network activity during navigation, these results outline a dynamic and distributed pattern of theta activity across the nodes of the navigation system (e.g. prefrontal cortex, posterior parietal cortex, parahippocampus), and highlight the utility of scalp recorded theta measures as potential indices of neural network function and hippocampal-parahippocampal physiology during navigation. We hope these findings warrant future investigations.

### Feedback processing

Consistent with previous work, the presentation of feedback stimuli in the T-maze elicited a large, focally distributed theta burst over the right temporal cortex (*17*). The topography and timing of this response are characteristic of RPT and indicate that the virtual reality T-maze paradigm is capable of eliciting this oscillatory response. Using a desktop version of the T-maze task, we demonstrated that RPT reflects a stimulus-induced partial phase reset (i.e. increase in power and enhanced phase consistency) of theta oscillations, and source localization, fMRI, and simultaneous EEG-fMRI data point to a neural generator in the right parahippocampal cortex (*17, 26, 39, 40*). In line with these observations, animal and computational work suggest that theta phase-coding and resetting are crucial during navigation as it sets the internal map of space encoded by the parahippocampal cortex (*6, 7, 10, 78, 79*). In order to prevent error accumulation of phase information during navigation, the phase of the theta rhythm may be reset to some predefined value (e.g. zero phase) by salient cues such as landmarks or rewards, a process thought to contribute to reward- and emotion-related spatial learning and memory(*6, 8*). Current thinking holds that this reset signal is provided by hippocampal place cells, which fire when the rodent enters the preferred field (or peak phase) of the place cell(*8, 78, 79*). More so, goal locations within a maze induces an accumulation of place fields and higher firing rates, which suggests that hippocampal place cells over-represent goal locations that generate emotional valence(*35*). Theta resets are also believed to be a mechanism for phase-locking hippocampal-parahippocampal activity to behaviorally relevant events and thereby may enhance cognitive processing (*7, 78, 80, 81*). By extension, we propose that the left and right goal locations within the T-maze were represented by it’s own place field.^i^ In particular, when the participant actively entered the goal location and received feedback, the phase of the parahippcampal theta oscillation was reset by the location-specific input from place cells, thereby concomitantly increasing theta phase coherence across trials. Further, the over-representation of goal locations by place cells (*35*) may have potentiated parahippocampal activity, thereby leading to an overall increase in regional spectral power. Accordingly, such stimulus-induced theta dynamics would be reflected in the EEG as enhanced theta phase consistency and spectral power, as we observed here with RPT power. In line with animal and computational work, we propose that RPT reflects a macroscopic proxy of the sum of parahippocampal theta activity, possibly the phase resetting of grid cells by place cells during feedback processing in the T-maze.

Next, we found that negative feedback relative to positive feedback yielded a significant increase in theta activity over frontal-midline electrodes, replicating the standard FMT effect(*37, 38*). At a behavioral level, participants exhibited a lose-switch strategy and walked faster on Win-stay trials^ii^, results suggesting that participants’ choices were influenced by the maze feedback. Over two decades of research using standard reinforcement learning paradigms (e.g. two-arm bandit task, gambling tasks, probabilistic reward tasks) have reliably demonstrated that FMT activities reflect the evaluation of negative and positive feedback for the purpose of the adaptive modification of behaviour (*37, 38, 64*). An accumulating body of evidence point to the anterior midcingulate cortex, as well as pre-supplemental motor area, as the source of FMT oscillations, and FMT power is thought to be modulated by a dopaminergic teaching signal tethered to prediction of reward outcomes during trial-and-error learning (i.e., reward predication error signals, RPEs)(*37, 38*). RPEs constitute the learning term in powerful reinforcement learning algorithms that indicate when events are “better” or “worse” than expected (*82*), and it is becoming increasing clear that positive and negative RPEs are encoded as phasic increases and decreases in the firing rate of midbrain dopamine neurons, respectively(*83*). FMT activities have also been shown to reflect a common computation used to identify and communicate the need for cognitive control, and subsequently organize prefrontal neuronal processes to implement top-down control across disparate brain regions(*37, 64*). By replicating the standard FMT response to reinforcers, in addition to the observed adaptive modification of behavior following feedback, we can infer the engagement of a reinforcement learning and control system during active navigation in this task.

In summary, **s**uccessful goal-directed navigation requires highly specialized neural representations that encode information about the location, direction, and speed of the navigating organism, as well as stimulus events, actions, and reinforcers for the purpose of optimizing behavior. Although substantial evidence from animal studies indicates that the theta rhythm plays a vital role in these neural representations during goal-directed navigation, they remain poorly understood in freely moving humans. In the present study, the multiplicity of human theta patterns observed during decision-making points, goal pursuits, and reward locations details how theta oscillations coordinate and support a diverse set of brain-wide neural assemblies and functions during goal-directed navigation. Foremost, measuring theta oscillatory activity from the scalp during active navigation allowed us to address our main objective: whether theta power increases with increases in speed, as shown previously in the rodent. This crucial finding opens a new door of investigative possibilities by which to integrate mobile-EEG measures of “real-life” goal-directed behavior with extensive animal, human, and computational work on spatial learning and memory based on Tolman’s seminal cognitive map theory.

## Materials and Methods

In this study, twenty-two young adults (20 right-handed [laterality index = 68], 9 male and 13 female, aged 18–29 years old [M = 21, SE =.61]) freely navigated a T-maze to find rewards (Fig. 1A). Participants were recruited from Rutgers University Department of Psychology subject pool using the SONA system. Each subject received course credit for their participation. Before the experiment, participants were screened for neurological symptoms and histories of neurological injuries (e.g., head trauma), and then asked to fill out the Edinburgh Handedness Inventory(*84*). After the experiment, participants filled out the Everyday Spatial Questionnaire. Ethical approval was obtained from the Institutional Review Board of the local university, and all participants provided written consent before the experiment.

In keeping with the classical design of the T-maze, this immersive virtual reality version consisted of a stem and 2 alleys extending at 90° angles out from a junction point and was located on a virtual enclosed landscape (20m × 20m) with an open ceiling exposed to a cloudy blue sky (Fig. 1; top panel). The virtual structure of the T-maze was enclosed inside the lab’s physical space of 2.13m by 2.13m room, with virtual meshed walls marking the boundaries. The T-maze was constructed using commercially available computer software (Unity version 2019.2, https://unity.com) and the virtual reality environment was provided through an HTC Vive head-mounted display system, which tracked participants’ head positions during navigation (HTC Corp., Taiwan). Continuous EEG was recorded with a mobile V-Amp amplifier from 16 actiCAP slim electrodes (Brain Products, Munich, Germany).

At the start of the experiment, a light beam marked the starting position of the T-maze, and the subjects had to step into that beam to start each trial. On each trial, participants walked down the stem of the maze until they reached a junction point, in which they were required to turn down the left or right alley and move towards a yellow orb floating at eye level at the end of the alley. The height of the icons was dynamically adjusted at the beginning of the experiment to match the subject’s eye-level. Once the participants were within 1.07 meters from the end of the alley, the floating yellow orb turned either green with a check mark (√) or red (x) for 1000 msec, signifying the alley they selected contained 5 cents (reward) or was empty (no-reward), respectively. Following the feedback, the maze would disappear, and participants were required to walk across an open field towards a purple beam of light. Once standing inside the beam of light and facing forward, the T-maze would re-appear, signifying the start the next trial. Participants were given 20 minutes to maximize their rewards. Unbeknownst to them, on each trial the type of feedback was selected at random (50% probability for each feedback type). At the end of the experiment, participants were informed about the probabilities and were given a $10 performance bonus.

The application contemporaneously communicated the subject’s position and the outcome of each trial by transmitting position values via a parallel port which took an integer from 0 to 255 and converted it to a voltage spike that was in turn marked by the EEG device. The rate of data updates was limited by the application’s running rate of 90 frames-per-second. Each signal was active for approximately 0.45 seconds, followed by transmitting a rest period of approximately seconds in order to allow for clear separation of the signals. However, the outcomes of each trial were recorded immediately, even if the aforementioned delay needed to be interrupted. The subject’s position was encoded as a 15 by 15 grid using integers 1 to 226, while outcomes were encoded using higher integers.

### Electrophysiological Data Recording

The electroencephalogram (EEG) data were collected using a 16-channel BrainVision actiCAP snap system (Brain Products GmbH, Munich, Germany) with 12 scalp electrode sites (Fp2, Fp1, Fz, Cz, FC5, FC6, Pz, Oz, P3, P4, P7, P8) and four external electrodes. One external electrode was placed on the right infraorbital region to record vertical eye movements (channel VEOG), and one was placed lateral to the outer canthus of the right eye to measure horizontal eye movements (channel RH). By convention, mastoid sites (M1 and M2) were collected to re-reference offline (see section below). EEG signals were recorded using Brain Vision Recorder software (Brain Products GmbH, Munich, Germany), online-referenced to channel FCz, a ground at AFz, and amplified using the portable V-Amp system (Brain Products GmbH, Munich, Germany). The sampling rate was set to 1000 Hz.

### Electrophysiological Data Reduction

Raw EEG recordings were analyzed offline using BrainVision Analyzer 2 (Brain Products GmbH, Munich, Germany). The first five trials were considered practice for each subject and were not included in the data analyses. We also excluded trials with response times (RTs) faster than 2.5% of the RT lower bound and slower than 2.5% of the RT upper bound to ensure the data quality. Raw EEG signals were filtered offline using a fourth-order digital Butterworth filter with a bandpass of .10-40 Hz. Activity at the online reference electrode FCz was recreated. Filtered signals were then subjected to ocular correction via independent component analysis (ICA). A mean slope algorithm was applied for blink detection, and an infomax-restricted algorithm was used for the ocular artifact correction. Channel Fp2 was used to detect vertical eye activity, and channel RH was used to detect horizontal eye activity. We then performed ICA correction on signals from 12 scalp electrodes (Fz, Cz, FC5, FC6, Pz, Oz, P3, P4, P7, P8, FCz, Fp1). Next, we divided the analysis stream into two pipelines: one for feedback-locked analyses and another for path analyses (i.e., from the starting point of one trial to the starting point of the next trial). For the feedback-locked analysis pipeline, signals were segmented into 5000 ms duration epochs spanning from −2500 ms to 2500 ms and time-locked to feedback onset. For the path analysis pipeline, signals were segmented into 25000 ms epochs time-locked to trial onset, spanning from −2500 ms to 22500 ms. Here, we used the prolonged epoch length for two reasons: (1) to ensure that the epoch was long enough to include the entire trial duration (i.e., from the start of one trial to the start of the next), and (2) to prevent the edge artifacts from time-frequency analyses. Following this, data were re-referenced using an average reference created from the following channels: FCz, Cz, FC5, FC6, Fz, Oz, P3, P4, P7, P8, and Pz. To note, by convention mastoid sites (M1 and M2) were collected to re-reference offline. However, these electrodes were removed from the dataset due to excessive noise and were not used in any of the analysis. Although mastoid references are commonly used in EEG research, future mobile virtual reality studies should avoid using this method as these channels tend to be contaminated by muscles involving in head rotation (e.g., sternocleidomastoid muscle).

For both pipelines, segmented data were then baseline-corrected using a mean voltage range from 200 ms to 0 ms preceding time 0. For feedback-locked segments, artifact rejection was conducted on the full segment with the following criteria: (1) a maximally allowed voltage step of 50 μV/ms, (2) a maximally allowed difference of values in intervals of 250 μV, and (3) lowest allowed activity values in intervals of 0.5 μV. For full-path segments, the search for artifacts was conducted within a customized window for each subject. The starting point of this customized window was −2500 ms relative to time 0. The endpoint of the window was the averaged RTs from the onset of one trial to the next across all trials plus 2500 ms. Due to the long epoch (25000 ms) used here, one segment often contained data from more than one trial—particularly for subjects with shorter RTs. By applying this customized window for each subject, we rejected epochs with artifacts that occurred within this interval of interest and preserved trials with artifacts that occurred outside of this interval (e.g., at the next trial) but not within. We added 2500 ms here to ensure that data points for convolution during time-frequency analyses were free from edge-artifacts to the greatest extent possible. On average, the duration of the customized window was 13014 ms (*SD* = 1304; min = 10789 ms; max = 15856 ms) across subjects included in the final data analyses (n = 22). After artifact rejection, bad channels (those with artifacts exceeding 5% of the data) were identified and interpolated using their four nearest neighbors’ signals for both pipelines (Hjorth, 1975). For subjects in the final analyses (n = 22), we interpolated data from one channel for four subjects (FC6: 1 subject; Oz: 1 subject; Cz: 1 subject; and FC5: 1 subject). All segmented data were written to individual MATLAB files for further processing using MATLAB software (MathWorks, Inc., 2019a). Out of 31 subjects whose EEGs were collected, data from 9 were excluded from final analyses due to multiple bad channels (n = 5), limited trial count (n = 2), extreme data outliers (n = 2), and failure to complete the experiment (n = 1).

### Time-frequency analyses

We conducted continuous wavelet transformation to decompose EEG oscillations into magnitude and phase information in the frequency range of 1 to 40 Hz for feedback-locked and full-path segments using a MATLAB program. For feedback-locked segments, the analysis was performed on four conditions: positive and negative feedback, rightward and leftward turns. For each condition, averaged evoked power was calculated by averaging the square of magnitude at each time point and frequency across trials. For feedback-locked segments, the analysis was performed on four conditions: positive and negative feedback, rightward and leftward turns. For each condition, averaged evoked power was calculated by averaging the square of magnitude at each time point and frequency across trials. To control for a potential difference in power spectrum before stimulus onset, we used a condition-average baseline of −300 to −150 ms pre-feedback onset averaging across all segments regardless of conditions for baseline normalization (*28*). For each subject, the power spectrum for the theta band (4-8 Hz) was averaged across all segments. We then identified the peak latency in the window of 0–600 ms post-stimulus for Fz and P8 (peak latency at Fz: 226 ms; peak latency at P8: 211 ms). The window for mean power extraction was then determined by +/− 25 ms around the peak latency for Fz and P8. We then used the window to extract mean amplitude for positive and negative feedback at Fz (window: 201–251 ms) and for leftward and rightward turns at P8 (window: 186–236 ms).

For the path analysis, we divided the segments into leftward and rightward turns based on their path choice for each subject. We also split the segments into fast and slow conditions based on the median RTs measured from trial onset to feedback across all segments for each subject. The averaged median RT was 3923 ms (*SD* = 624; min = 2893; max = 5473) across 22 subjects. The segments were then subjected to continuous wavelet transformation for each condition. After the transformation, a critical challenge for creating an average power spectrum was that the timing of event triggers marking the turn and feedback location in time relative to time 0 (i.e., trial onset) varied across segments. Such variation made it challenging to obtain a robust averaged power spectrum using the conventional averaging approach. Therefore, we applied a data binning strategy used in animal studies to examine neurophysiology in freely moving rats to address timing variation across trials (e.g., Kyriazi, Headley, & Pare, 2020).

To apply the binning strategy, we divided each segment into two sections (Stem and Turn) according to the triggers marking participants’ movement trajectories in the T-maze. The Stem section was defined as the period between trial onset and the intersection of the T-maze. The Turn section was defined as the period between the junction of the T-maze and feedback onset. We then binned the power spectrum into 60 bins for each defined maze section using the *histcounts* function written in MATLAB (Mathworks Inc., Natick, MA). Specifically, for a given section, the program divided the interval in milliseconds into approximately equally spaced bins and defined the bin edges (i.e., the starting point and the endpoint in milliseconds). We then averaged the total power across the time points in milliseconds within each bin. For example, for a given trial, the duration of the Stem section was 1500 ms, indicating that the width of each bin is 25 ms. We would then average the total power across 1-25 ms to get the total power for bin 1; average the total power across 26-50 ms to get the total power for bin 2; average the total power across 51-75 ms to get the total power for bin 3, etc. We did this for each frequency in every trial. We then averaged single-trial binned total power across segments for each condition to obtain the averaged binned total power for each subject. For both the path analyses, the averaged binned total power was then baseline normalized using a condition-average baseline (i.e. all conditions averaged together) in the period of −1000 ms to −100 ms before the trial onset. Across these 22 subjects, the averaged milliseconds per bin were 42 ms for the Stem section and 27 ms for the Turn section. The mean power was extracted across all channels for delta, theta, and alpha bands for the following sections (Figure 3): (1) S1a: Stem section – first half (Bins 1-30: first half of trajectory from start location to junction point); (2) S1b: Stem section – second half (Bins 31-60: second half of trajectory from start location to junction point); (3) S2a: Turn section – first half (Bins 61-90: first half of trajectory from junction point to left or right feedback location); and (4) S2b: Turn section – second half (Bins 91-120: second half of trajectory from junction point to left or right feedback location. To note, because of the inter-trial and inter-subject variation in return strategies (e.g. turn counter-clockwise vs clockwise to return to start location; walk forward vs backwards to start location – information was not recorded), we did not include an analysis of the return segment of the task and leave this for future investigations. All statistical analyses were performed using SPSS 24.0 for Windows (IBM SPSS Statistics, IBM Corporation).

## Acknowledgments

We are grateful to D.H., M.X.C., J.C., and J.R. for consultation on the time-frequency analysis and to the research assistants of the Laboratory for Cognitive Neuroimaging and Stimulation for help with data collection.

## Author contributions

Conceptualization: TEB, OL
Methodology: TEB, OL, ML
Investigation: ML, NB, TEB
Visualization: TEB
Supervision: TEB
Writing—original draft: TEB, ML
Writing—review & editing: OL

## Data and materials availability

The data that support the findings of this study are available from the corresponding author (TEB) on request.

## Footnotes

i. This idea may explain why we failed to replicate the rightward turning bias on RPT power and latency (phase) observed in our previous 2D T-maze tasks(*17*). For instance, during active navigation, if the two goal locations were represented by their own place fields, and the feedback-induced reset occurred at the center of each place field, then the resulting RPT phase and power would be identical between the two goal locations. By contrast, if the two goal locations in the 2D version of T-maze task were only represented by one place field — since subjects were only sitting in one physical location and pressing buttons to move between different spatial locations digitally drawn on the screen — it is possible that the left and right goal location were represented by different phase positions along the theta cycle of a single place field. If true, one might expect to see commensurate differences in RPT power and phase between left and right goal locations following phase reset, as we observed previously(*17*).

ii. Its interesting to note that we failed to replicate the standard win-stay behavior, a heuristic learning strategy used to model learning in decision situations and has been applied towards theory development in psychology, game theory, statistics, economics, and machine learning (*38, 85, 86*). In particular, when subjects are simply pressing buttons to make decisions on a computer, this win-stay pattern emerges (*87, 88*), but when subjects are required to move their entire body to make decisions, this pattern disappears. While this is a surprising result and needs to be explored further, it is our best guess that the win-stay and win-shift decisions during active navigation reflects an increase in strategy exploration (testing win-shift behavior more often) or there are differences in the computations between active navigation (i.e. calculating the physical and cognitive energy needed to navigate our bodies towards a goal), and simple button presses.

